# Aberrant medial prefrontal cortex activity and flexible behavior in the TgF344-AD rat model of Alzheimer’s disease

**DOI:** 10.1101/2025.07.10.664159

**Authors:** T. Joseph Sloand, Benjamin P. Dunham, Mark Niedringhaus, Elizabeth A. West

**Affiliations:** Department of Neuroscience, School of Osteopathic Medicine and School of Translational Biomedical Engineering and Sciences, Virtua Health College of Medicine and Life Sciences, Rowan University, Stratford, NJ 08084

## Abstract

Cognitive deficits, including deficits in the ability to shift behavior following negative consequences, often precede the accumulation of canonical neuropathological markers (Aβ plaques and tauopathy) and severe dementia in Alzheimer’s disease (AD) patients. The Tg-F344-AD rat model exhibits age-dependent AD pathology and memory deficits that recapitulate AD, However, it is unknown how medial prefrontal cortex activity is altered in awake and behaving AD rats during learning and/or flexible behavior. Here we determine the ability of in 6-7month-old TgF344-AD rats to learn reward predictive cues and to shift behavior away from reward-predictive cues following outcome devaluation while recording mPFC neurons. Specifically, AD rats (n=17) and wild-type littermates (n=17) were presented with two distinct cues as conditioned stimuli (CS+) predicting distinct outcomes. A conditioned taste aversion to one outcome was induced, after which the rats were tested post-devaluation to evaluate their ability to avoid the CS+ associated with the devalued outcome. We found a loss of motivated behavior during learning and a loss of flexible behavior during testing in 6-7-month-old AD rats relative to WT littermate controls. In addition, there was differential aberrant mPFC encoding of cue-outcome associations in AD rats during conditioning and following outcome devaluation. Specifically, AD animals show fewer neurons during conditioning that encode both the cue and the outcome than WT animals. Also, AD animals also showed a greater proportion of neurons that exhibited an excited response to reward predictive cues post-outcome devaluation. Together, these data contribute to our understanding of alterations in mPFC that may underline prodromal AD behavioral deficits to inform future treatments.

## Introduction

Alzheimer’s disease (AD) is a progressive neurodegenerative disorder and one of the leading causes of dementia in older adults, and AD diagnoses are expected to dramatically increase (Report, 2024). While the neuropathology of AD is often characterized in the context of hippocampal dysfunction (Visser et al., 2002; Berron et al., 2020), altered activity in prefrontal cortical (Kaufman et al., 2012; Bertoux et al., 2015) may also play a role in the cognitive deficits of early AD (Fowler et al., 2022). The cognitive deficits of AD, including impairments in executive function, attention, and decision making are likely apparent before the accumulation of classical neuropathological markers like Aβ, tau, and gross neuronal loss (Koenig et al., 2005; Hamm et al., 2015; de Siqueira et al., 2017), however, given that age also influences these executive functions it is difficult to diagnosis AD prior to classic memory deficits. Therefore, utilizing an animal model that recapitulates classic pathology and memory deficits holds great translational value to determine how behavioral changes occur prior to pathology.

Flexible behavior, defined as the ability to shift behavior following changes in outcome value, is disrupted in both human populations (Freedman and Oscar-Berman, 1989) and animal models (Rorabaugh et al., 2017) of early AD, and these deficits likely lead to dysfunctional decision making (Creese and Ismail, 2022). Indeed, AD-induced neuropathological changes occur in the prefrontal cortex (Igarashi et al., 2011; Beagle et al., 2020). In the rat, the medial prefrontal cortex (mPFC) is involved in altering behavior to predicted outcome values (O’Doherty, 2011; McNamee et al., 2015; West et al., 2021; Niedringhaus and West, 2022). Flexible behavior can be measured using the outcome devaluation task, where performance on the task depends on a functional prelimbic subregion of the mPFC during learning (West et al., 2021). We have recently shown that mPFC neural activity to a cue that predicts the devalued outcome correlates with the ability to shift behavior post-outcome devaluation (Niedringhaus and West, 2022).

The TgF344-AD model, a rat model of AD exhibiting age-dependent development of pathological signatures and memory deficits closely recapitulates human AD (Cohen et al., 2013). The TgF344-AD rats show deficits in the ability to shift behavior in a water-maze reversal learning by 6 months of age (Rorabaugh et al., 2017) prior to any hippocampal pathology or classical AD neuropathology (Cohen et al., 2013). Moreover, AD rats show dampened oscillatory dynamics (*in vivo* electrophysiology in anesthetized rats) within the rodent mPFC at the relatively early time point (Bazzigaluppi et al., 2018). Critically, in both humans (Koenig et al., 2005; Hamm et al., 2015) and rats (Bazzigaluppi et al., 2018), neurophysiological changes in the prefrontal cortex predate not only the histopathological changes associated with AD (i.e., Aβ and tau accumulation) but also the learning and memory deficits resulting from hippocampal pathology later in the disease progression (Cohen et al., 2013). However, it is unknown how neural activity in the mPFC is altered in AD rats throughout learning and subsequent flexible behavior. Further, to date, there have been no electrophysiological studies in *awake and behaving* AD rats. Here, we used *in vivo* electrophysiological recordings in awake, behaving rats to examine the mPFC encoding of reward predictive cues before and following outcome devaluation in TgF344-AD rats and age-matched littermate controls.

## Materials and Methods

### Subjects

34 heterozygous TgF344-AD rats and age-matched (6-7 months old) WT littermate controls (Cohen et al 2013) were used from our in-house colony (n=10 male WT, n=7 female WT, n=8 male TgF344-AD, n=8 female TgF344-AD). To establish the colony, male TgF344-AD rats (Rat Resource and Research Center, Columbia, MO) were bred with purchased (Envigo) F344 females, and pups were ear punched and genotyped using PCR. All rats were housed individually and maintained on a biphasic light cycle (lights off at 9:00 PM, lights on at 9:00 AM). During all behavioral procedures, rats were food-regulated to no less than 85% of their free-fed body weight. All animal procedures were approved by the Rowan University Institutional Animal Care and Use Committee (IACUC).

### Surgical Procedures

Rats were anesthetized with a mixture of ketamine hydrochloride (100 mg/kg) and xylazine (10 mg/kg, i.p.), and microwire recording arrays were bilaterally implanted into the mPFC (AP: +2.5-2.8 mm, ML: ±0.5-0.6 mm, DV: −4 --5 mm).

### Behavioral Task

Rats were trained on a reinforcer devaluation task that consisted of 3 phases: Pavlovian conditioning, devaluation, and a post-devaluation test as shown in Figure 1A.

**Figure 1.**
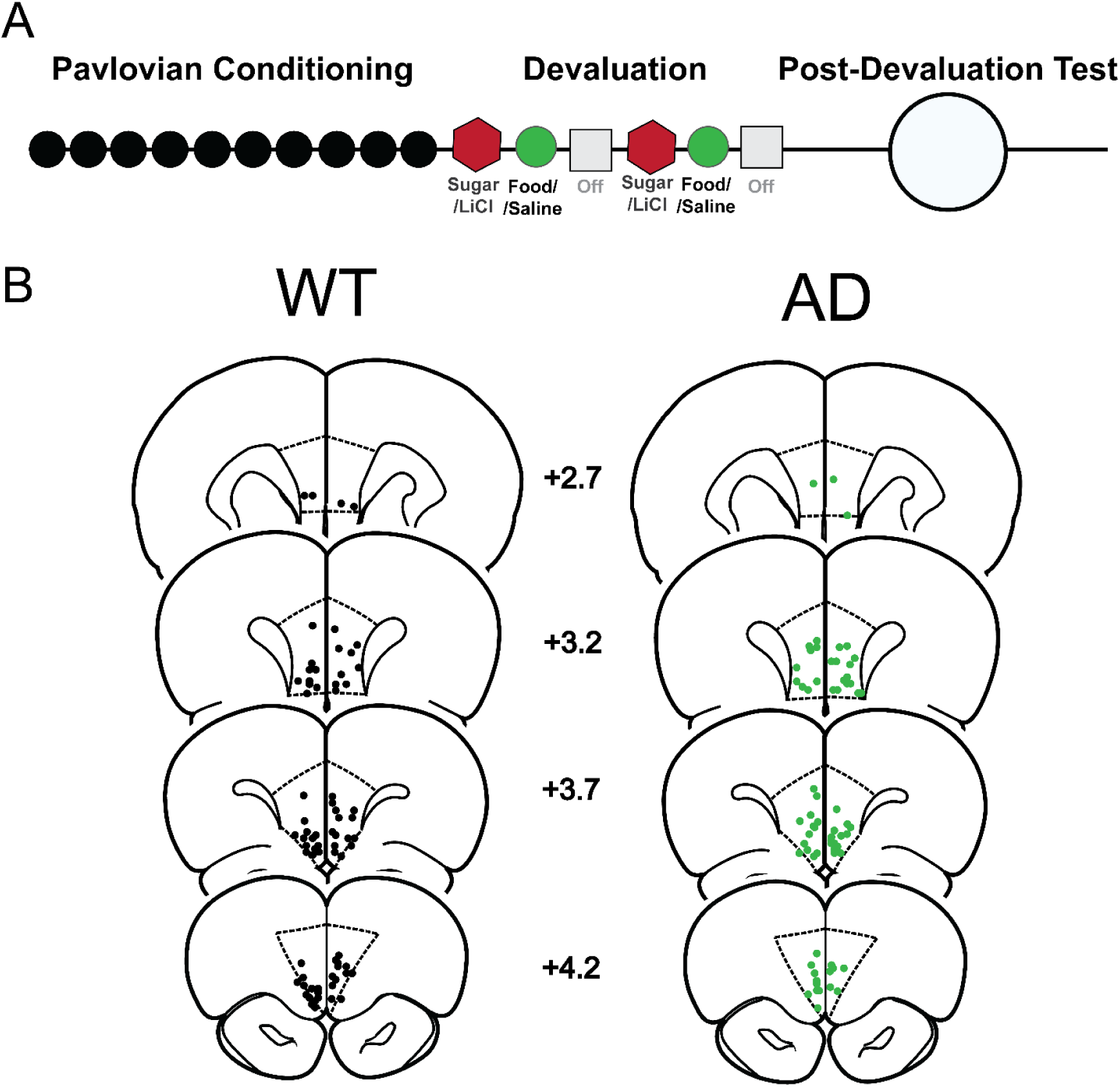
Schematics of behavioral training and testing timeline and array placements. A) Behavioral timeline of the experimental approach, including 10 days of Pavlovian conditioning (black dots), 6 days of the devaluation procedure (red hexagon-sugar paired with LiCl, green circle, food paired with saline, grey square-day off), and 1 post-devaluation test day (grey circle). B) Microwire placements in the medial prefrontal cortex in WT (left) and AD (right) rats.

#### Pavlovian Conditioning

Rats underwent 10 days of Pavlovian conditioning where they were presented with two distinct cue patterns as conditioned stimuli (CS+, 10 s each, solid or flashing) that each predicted a sucrose or grain pellet (45 mg, bio-serv) counterbalanced and two other cue patterns (solid or flashing, in a different spatial location) that did not predict a reward (CS−); 10 trials each for 10 daily sessions (West et al., 2021; Niedringhaus and West, 2022).

#### Devaluation

Rats were habituated for three days in a standard empty rat cage. Following habituation, rats were given free access in this empty cage to sucrose pellets for 30 minutes immediately before receiving an injection of LiCl (i.p., 0.3 M, 7.5 ml/kg) to induce a conditioned taste aversion. The next day, rats were given free access to grain pellets for 30 minutes immediately before receiving a saline injection (i.p., 7.5 ml/kg). At least 48 hours later, the same two-day devaluation procedure was repeated (West et al., 2021; Niedringhaus and West, 2022). To prevent any interaction with rats’ normal home cage chow, sucrose pellets were exclusively devalued in this study.

#### Post-Devaluation Test

At least 48 hours after the final devaluation session, rats underwent a single session of the Pavlovian conditioning procedure as described above but with no rewards delivered (under extinction). On the following day, rats were given free access to both rewards for 30 minutes in a food choice consummatory to confirm successful devaluation (West et al., 2021; Niedringhaus and West, 2022).

### In Vivo Electrophysiology

mPFC neural activity was recorded during every conditioning and test session using a neurophysiological system (Omniplex, Plexon Inc., Dallas TX). Continuous signal from each electrode was virtually referenced (PlexControl, Plexon Inc., Dallas TX) and was high-pass filtered (300 Hz) to identify individual spike events. Cell recognition and sorting was finalized using the Offline Sorter software (Plexon Inc., Dallas TX). Waveform and spontaneous firing rates were examined to identify putative pyramidal neurons in the mPFC (Moorman and Aston-Jones, 2015). A separate computer processed operant chamber events (Med Associates, Inc.) and sent timestamps of the events to Omniplex such that neural data could be time-locked to behavioral events.

### Data Analysis

#### Behavior

Videos were scored using Cineplex software by an observer who was blind to the experimental treatment. The amount of time spent in the food cup into the food cup during each cue presentation was measured for each animal on Day 1 and Day 10 of conditioning and analyzed using a three-way mixed ANOVA with Day (1 vs. 10) and cue type (both CS+ vs. both CS−) as within subject factors and genotype (AD vs. WT) as a between subject factor. In addition, we analyzed the day 10 conditioning data as a two-way mixed ANOVA with cue type (CS+ vs CS−) as within-subject and genotype (AD vs WT) as between-subject. Based on the % of time spent in the CS+ compared to the CS−, we calculated a conditioning index using the following formula: [(CS+ - CS−)/(CS+ + CS−) (West et al., 2021). For the post-devaluation test day, we analyzed these data using a two-way repeated measures ANOVA with the following factors: genotype (WT vs. AD) and devaluation status (nondevalued, ND, vs. devalued, D), including the first trial of each cue only (Sood and Richard, 2023; Pickens et al., 2024) to avoid the influence of extinction testing. Based on the % of time spent in the nondevalued (ND) condition compared to the devalued (D) condition, we calculated a devaluation index [DI (West et al., 2012; West et al., 2013; West and Carelli, 2016; West et al., 2021; Niedringhaus and West, 2022)] on the first trial of the post-devaluation test session using the following formula: (CS^+^ predicting nondevalued (ND) outcome - CS^+^ predicting devalued (D) outcome)/ (CS^+^ predicting nondevalued (ND) outcome + CS^+^ predicting devalued (D) outcome)/ or (ND-D)/(ND+D). To normalize this to Day 10 preferences, we used the same formula as above on Day 10 and subtracted it from the index on the test day (West et al., 2021). The amount of food consumed (g) of both outcomes (grain and sucrose pellets, post-devaluation) was measured for the post-test food choice. Consumption during the food choice test was analyzed using a two-way ANOVA with genotype (AD vs. WT) and devaluation status (ND vs. D) as factors. We used SPSS software (IBM) for behavioral analyses.

#### Electrophysiology

Data were analyzed blind to treatment. Peri-event histograms (200-ms bins, 30s total) were used to analyze neuronal activity relative to task events (cue on) on either the last day of conditioning (i.e., Day 10) or the post-devaluation test session. Putative pyramidal neurons were identified based on firing rate and waveform width (Moorman and Aston-Jones, 2015). Individual neurons were classified as phasic if, during the cue period, the firing rate was greater than (excitation) or less than (inhibition) the baseline period as measured by a student’s paired t-test across baseline bins and event period bins (either cue period or outcome period). Units that exhibited both excited and inhibited profiles during the same cue period were classified by the response that was most proximal to the event. We analyzed the proportion of cells classified as nonphasic, excited, or inhibited using the chi-square test on Day 10 (CS+ vs CS−) and the Post-Devaluation Test Day. Next, for each cell, we calculated a z-score [(x – μ) / σ] for analysis examining z-score firing rates across groups. Here, we only included neurons that were monophasic (i.e., only an excited or an inhibited response) but not biphasic (6 neurons were excited and inhibited to the CS+ in the WT group, and 1 neuron was excited and inhibited to the CS+ in the AD group) for comparing the normalized firing rate of excited and inhibited neurons between WT and AD rats. Specifically, for all excited and inhibited neurons (monophasic), we calculated the area under the curve (AUC) during the cue period and performed two-way ANOVAs for group and cue type (CS+ vs CS− or nondevalued vs devalued). We performed linear regression for performance based on the devaluation index (see behavior analysis) and phasic neural encoding (% of phasic neurons in animals in which we recorded at least 4 neurons in the session). We used GraphPad Prism for chi-square, AUC, and linear regression analyses.

### Histology

Animals were deeply anesthetized using vaporized isoflurane and transcardially perfused with physiological saline and 4% paraformaldehyde. After post-fixing and freezing, 40-μm coronal brain sections were examined to verify microwire array placement. Placement of the microwire arrays in WT and AD rats are shown in Figure 1B.

## Results

Transgenic AD rats show diminished approach behavior towards reward predictive cues but successfully discriminate between rewarded and unrewarded cues (CS+ vs CS−). After 10 days of Pavlovian conditioning, both groups of rats spent more time in the food cup to CS+ compared to the CS−, but AD rats spent less time in the food cup in the reward predictive cues (CS+ and CS) compared to WT as shown in Fig. 2A-B. Specifically, a three-way mixed analysis of variance (ANOVA) revealed a main effect of day (F_(1,32)_=138.4, p<0.001), cue type (F_(1,32)_=88.9, p<0.001), and genotype (F_(1,32)_=4.5, p=0.04). In addition, there was a significant interaction for day x genotype (F_(1,32)_=4.4, p=0.04), day x cue type (F_(1,32)_=133.4, p<0.001), genotype x cue type (F_(1,32)_=7.1, p=0.01), and critically a day x cue type x genotype interaction (F_(1,32)_=7.3, p=0.01). A post-hoc Bonferroni test corrected for multiple comparisons (one-factor difference compared) revealed both groups increased time spent in the food cup from Day 1 to Day 10 for both cue types [WT: CS+ (F_(1,32)_=145.4, p<0.001), CS− (F_(1,32)_=22.5, p<0.001); AD: CS+ (F_(1,32)_=65.4, p<0.001), CS− (F_(1,32)_=13.4, p<0.001] and that WT rats spent more time in the food cup on Day 10 compared to AD rats during the CS+ (F_(1,32)_=9.5, p=0.004), but not to the CS− (F_(1,32)_=1.0, p=0.32). A two-way mixed ANOVA revealed a significant main effect of genotype (F_(1,32)_=4.9, p=0.03), cue type (F_(1,32)_=122.5, p<0.0001), and genotype x cue type interaction (F_(1,32)_=7.8, p=0.009) on the percent of time spent in the food cup on Day 10 of conditioning (Fig. 2B). Critically, AD rats spent significantly less time in the food cup during the CS+ presentation than WT rats as revealed by Bonferroni post-hocs corrected for multiple comparisons (CS+; F_(1,32)_=9.5, p=0.004), CS−; (F_(1,32)_=1.0, p=0.32), even though both groups spent more time in the foodcup to the CS+ relative to the CS− [WT; (F_(1,32)_=95.1, p<0.001); AD: (F_(1,32)_=33.3, p<0.001]. To determine if the effect was due to differences in the ability to discriminate between the CS+ vs CS− or a potential general decrease in motivation towards the CS+, we calculated a conditioning index (cite, CS+ - CS−/ CS+ + CS−) in which 1 would represent complete discrimination (i.e., only responding towards the CS+, but not responding to the CS−) and 0 would represent equal responding to both the CS+ and CS−. We found no difference between the WT and AD transgenic rats on the conditioning index as shown by an unpaired t-test (t_32_=1.5, p=0.14), and both groups demonstrated successful discrimination as measured by a one samples t-test (WT: t_16_=9.2, p<0.0001, AD: t_16_=5.7, p<0.0001) compared to the hypothetical value of 0 (i.e., no discrimination) as shown in Figure 2C.

**Figure 2.**
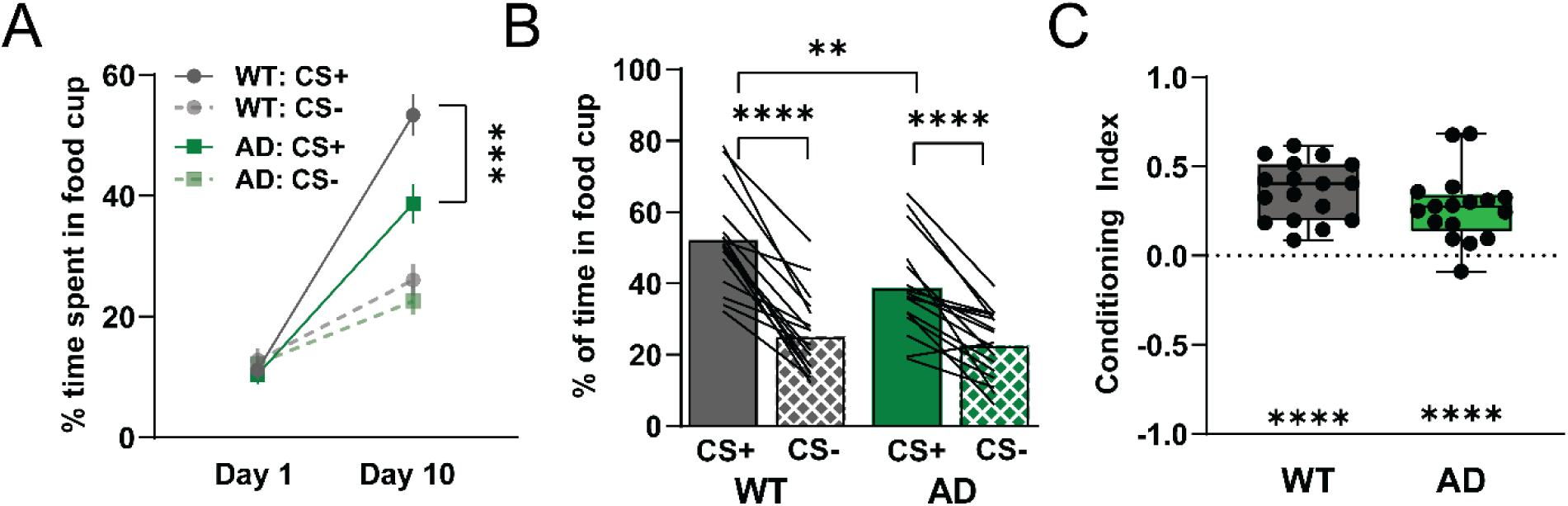
Transgenic AD rats show diminished approach behavior towards reward predictive cues but successfully discriminate between reward and unrewarded cues (CS+ vs CS−). A) After 10 days of Pavlovian conditioning, both groups of rats spent more time in the food cup to CS+ compared to the CS−, but AD rats spent less time in the food cup in the reward predictive cues (CS+ and CS) compared to WT [main effect of day (F_(1,32)_=138.4, p<0.0001), cue type (F_(1,32)_=88.9, p<0.0001), and genotype (F_(1,32)_=4.5, p=0.04) and a day x cue type x genotype interaction (F_(1,32)_=7.3, p=0.01)]. A post-hoc Bonferroni test corrected for multiple comparisons revealed that WT rats spent more time in the food cup on Day 10 compared to AD rats during the CS+ (*** represents p<0.001), but not to the CS−. B) On Day 10 of conditioning, AD rats spent significantly less time in the food cup specifically during the CS+ presentation than WT rats [main effect of genotype (F_(1,32)_=4.9, p=0.03), cue type (F_(1,32)_=122.5, p<0.0001), and genotype x cue type interaction (F_(1,32)_=7.8, p=0.009); with Bonferroni post-hocs comparing AD rats and WT rats (** represents p<0.01] even though both groups of rats spent more time in the food cup during the CS+ relative to CS− (****p<0.0001). C) There was no difference in the ability to discriminate between CS+ and CS− between the WT and AD transgenic rats as measured by a conditioning index (CS+ - CS−/CS+ + CS−) as shown by a paired t-test (t_32_=1.5, p=0.14), and both groups demonstrated successful discrimination as measured by a one samples t-test (WT: t_16_=9.2, p<0.0001, ****, AD: t_16_=5.7, p<0.0001, ****) compared to the hypothetical value of 0 (i.e., no discrimination).

On Day 10 of conditioning, we recorded from 275 neurons (AD: 118; WT: 157) in the medial prefrontal cortex (PrL and IL subregions) on the final day of conditioning (Figure 1A). Figure 1B shows the distribution of microwire array placements in WT and AD rats. We found that distinct population showed an increase (excited, “EXC”) or a decrease (inhibited, “INH”) to the cue presentation and thus classified as phasic to either the CS+ or the CS−, while cells that do not decrease or increase in firing to the were classified as nonphasic (“NP”) as shown in Figure 3A-B. We did observe a significant difference between the proportion of nonphasic, excited, and inhibited cells to the cues (CS+/CS−) on Day 10 between AD and WT animals as shown by a χ^2^ analysis (χ^2^=16.8, df=6, p=0.01) as shown in Figure 3A-B. We found a difference between CS+ vs CS− in both the WT (χ^2^=6.2, df=2, p=0.04). and the AD group (χ^2^=6.3, df=2, p=0.04). We did not observe a difference between the CS− (χ^2^=3.3, df=2, p=0.19) or CS+ (χ^2^=1.5, df=2, p=0.46 between AD and WT animals. Next, we examined the proportion of only phasic neurons to determine the differences between AD and WT in neural encoding. We found that on Day 10 there was no difference in the proportion of excited vs inhibited classified neurons as shown in Figure 4C (χ^2^=0.72, df=3, p=0.86). Finally, we found a difference between neural encoding between the CS+ vs CS−, but no differences between in AD and WT rats as shown in Figure 3D. Specifically, we calculated an AUC for the cue duration (either CS+ or CS−) for each neuron and a two-way ANOVA revealed a main effect of cue type (CS+ vs CS−) but no main effect of group (AD vs WT) or interaction for neurons classified as excited (group: F_(1,150)_=0.39, p=0.53; cue type: F_(1,150)_=4.6, p=0.03; interaction F_(1,151)_=0.005, p=0.94) and inhibited (group: F_(1,89)_=3.4, p=0.07; cue type: F_(1,89)_=14.6, p<0.001; interaction F_(1,89)_=0.88, p=0.35).

**Figure 3.**
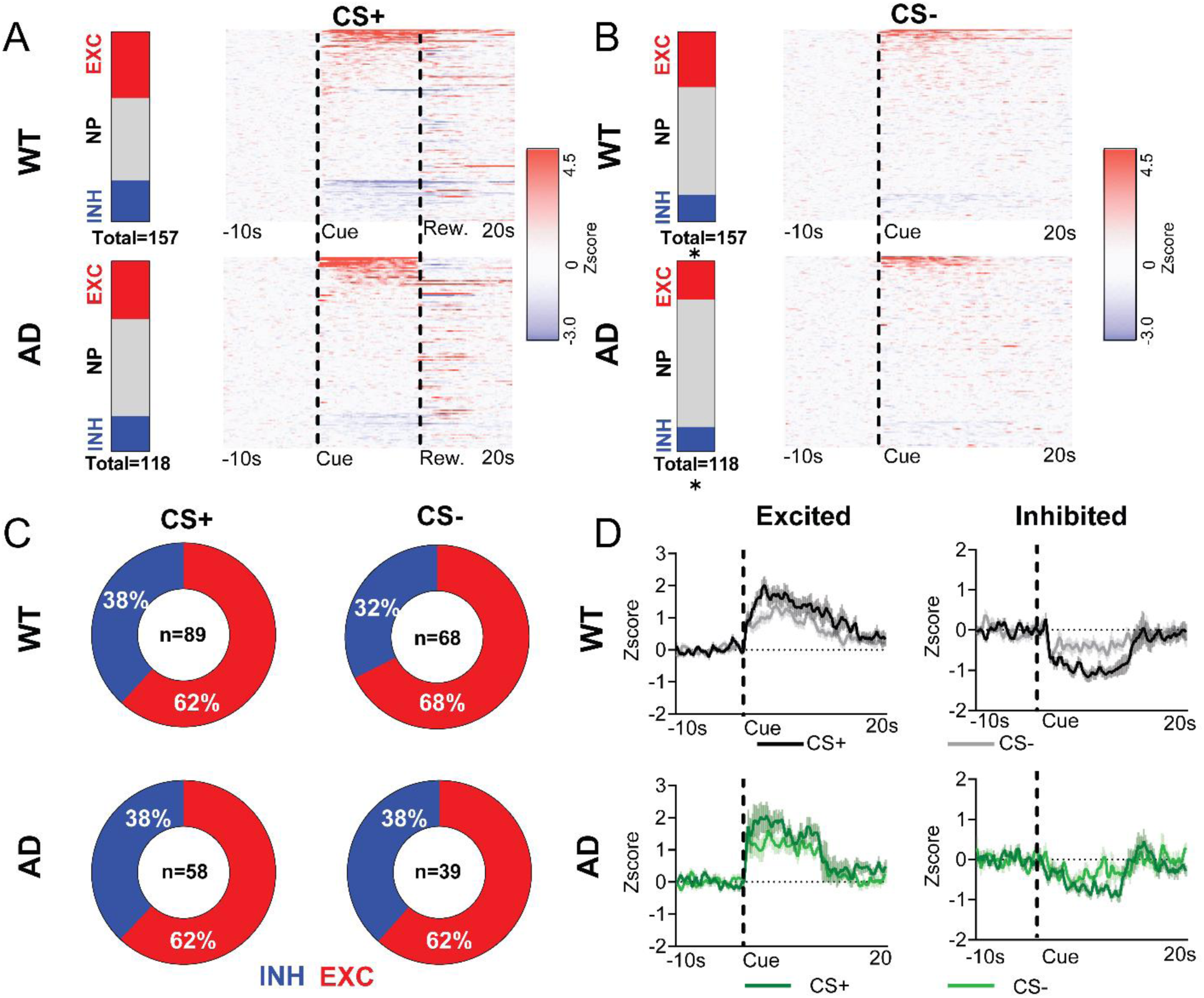
Both AD and WT rats show preferential neural encoding to the CS+ relative to CS−. A) Proportion of excited, inhibited and nonphasic neurons to reward predictive cue (CS+) in WT and AD and corresponding heatmap of all neurons recorded. B) Proportion of excited, inhibited and nonphasic neurons to the CS− (i.e., no predictive reward) in WT and AD and corresponding heatmap of all neurons recorded showing a significant difference between the proportion of neuron classification between CS+ and CS− for AD and WT animals (* p<0.05). C) Piecharts representing the proportion of phasic neurons classified as excited (EXC) or inhibited (INH) show no difference in the proportion between AD and WT to CS+ vs CS− D) The average firing (zscore) across all neurons classified as either excited (left) and inhibited (right) in WT and AD rats with AUC during the cue period (10s) difference between CS+ and CS−, but no difference between WT and AD rats.

**Figure 4.**
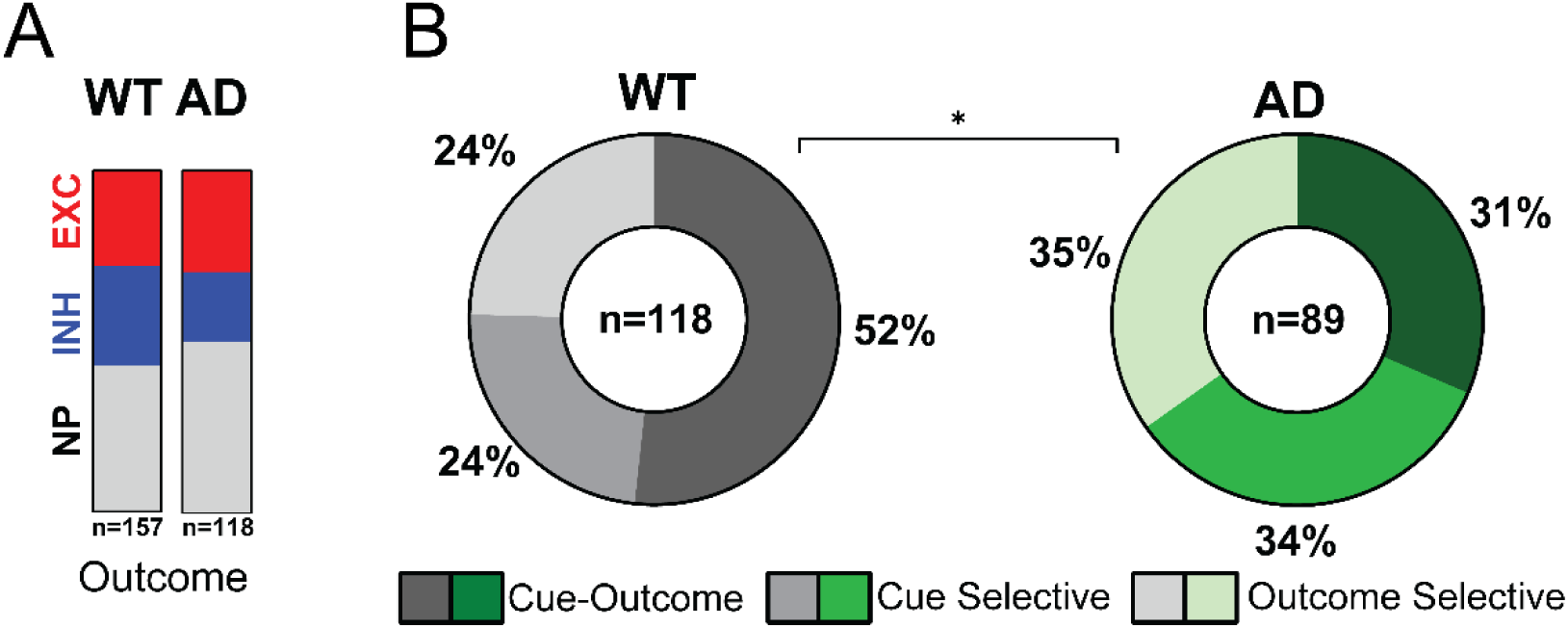
AD and WT show differential cue-outcome encoding on the last day of conditioning. A) There is no difference between the proportion of neurons that were excited, inhibited, or nonphasic to the outcome in AD and WT animals. B) WT animals show a significant increase in the proportion of neurons that encode both the cue and the outcome compared to AD animals that show equal number of neurons that were classified as cue-outcome (both cue and outcome), cue selective (cue only), or outcome selective (outcome only) as shown by chi-square (χ^2^=4.9, df=1, * p=0.03).

We did not observe a difference in the proportion of nonphasic, excited, or inhibited cells that responded to the outcome on Day 10 (χ^2^=3.0, df=2, p=0.23) between WT and AD animals as shown in Figure 4A. However, when we classified phasic neurons into whether the neurons responded to the cue and outcome (Cue-Outcome), cue only (Cue Selective), or outcome only (Outcome Selective), we found that there was a different proportion of classification between WT and AD (χ^2^=8.5, df=2, p=0.01). Specifically, WT animals show predominantly cue-outcome phasic neurons (52%), thus neurons that respond to both the cue and the outcome. In contrast, AD animals showed an equal proportion of mPFC neurons that were cue-outcome, cue-selective, and outcome selective, as shown in Figure 4B.

On the post-devaluation test session, AD rats showed diminished motivated behavior and impaired ability to shift behavior away from the cue predicting the devalued outcome. On post-devaluation test day, a two-way mixed ANOVA revealed a significant main effect of genotype (F_(1,32)_=8.0, p=0.008, but not of devaluation status (F_(1,32)_=3.0, p=0.091) or devaluation status x genotype interaction (F_(1,32)_=2.05, p=0.16). Planned comparisons separated by one factor (Bonferroni corrected for multiple comparisons) revealed that WT rats spent significantly less time in the food cup during the cue that predicted the devalued outcome compared to the cue that predicted the nondevalued outcome (F_(1,32)_=5.0, p=0.032), while there was no difference between the time spent in the food cup during the cue that predicted the two outcomes (F_(1,32)_=0..05, p=0.82) in AD rats as shown in Figure 5A. Furthermore, we found a significant difference between WT and AD rats in the amount of time spent in the foodcup during the cue associated with the nondevalued outcome (F_(1,32)_=12.4, p=0.001), but not the devalued cue (F_(1,32)_=1.5, p=0.22). Given that we observed significantly diminished responding in general across conditioning in the AD rats compared to WT rats (Figure 2A-B), we examined the degree of responding to both cues on the Post-Devaluation Test Day relative to the degree of responding on Day 10 for the respective cues (% time spent on test day - % time spent on Day 10) to control for general diminished motivated behavior. We found that WT animals significantly decreased their responses to cues that predicted the devalued outcome compared to the nondevalued outcome, while AD animals did not, as revealed by a mixed two-way ANOVA with a significant devaluation status x genotype interaction (F_(1,32)_=6.2, p=0.02), and no main effect of devaluation status (F_(1,32)_=3.1, p=0.09) or genotype (F_(1,32)_=0.15, p=0.70). Bonferroni post-hoc revealed a significant difference between the change in responding in the devalued condition compared to the nondevalued in WT (F_(1,32)_=9.0, p=0.005), but not in AD rats (F_(1,32)_=0.28, p=0.28). We also calculated a devaluation index [ND - D/ ND + D] in which 1 would represent maximally flexible behavior (i.e., only responding towards the nondevalued cue and avoiding the devalued cue) and 0 would represent equal responding to both cues and impaired ability to flexibly adjust behavior following devaluation. We found no difference between the WT and AD transgenic rats on the devaluation index as shown by a paired t-test (t_32_=0.92, p=0.37), however only WT rats showed successful flexible behavior as WT showed a significant positive value for the devaluation index compared to the hypothetical value of 0, while AD rats’ performance did not differ from 0 indicative of impaired flexible behavior as shown by a one-sample t-test (WT: t_16_=3.2, p=0.005; AD: t_16_=0.32, p=0.33, Figure 4C). Finally, to confirm that there were no differences in the ability of the rats to devalue the outcome more generally after pairing it with LiCl, we ran a consummatory food choice test in which rats had free access to both the nondevalued and devalued outcome. WT and AD rats show successful devaluation of the outcome, but AD rats also show a decrease in the general consumption of the nondevalued outcome as well (Fig. 4D) as revealed by a two-way mixed ANOVA with a main effect of devaluation status (F_(1,32)_=126.6, p<0.0001), main effect of genotype (F_(1,32)_=4.5, p=0.04), and interaction between genotype x devaluation status (F_(1,32)_=4.6, p=0.04). Bonferroni post-hocs corrected for multiple comparisons showed that both WT and AD rats ate significantly more of the nondevalued outcome than the devalued outcome (WT: F_(1,32)_=89.8, p<0.0001, AD: F_(1,32)_=41.4, p<0.0001). In addition, AD rats ate significantly less of the nondevalued outcome than WT rats (F_(1,32)_=4.5, p=0.04). There was no difference in the amount of devalued outcome consumed (F_(1,32)_=1.0, p=0.32). Together, our behavioral findings reveal two behavioral phenotypes in AD rats, one a general loss of motivation as well as an inability to change behavior after a change in devaluation.

**Figure 5.**
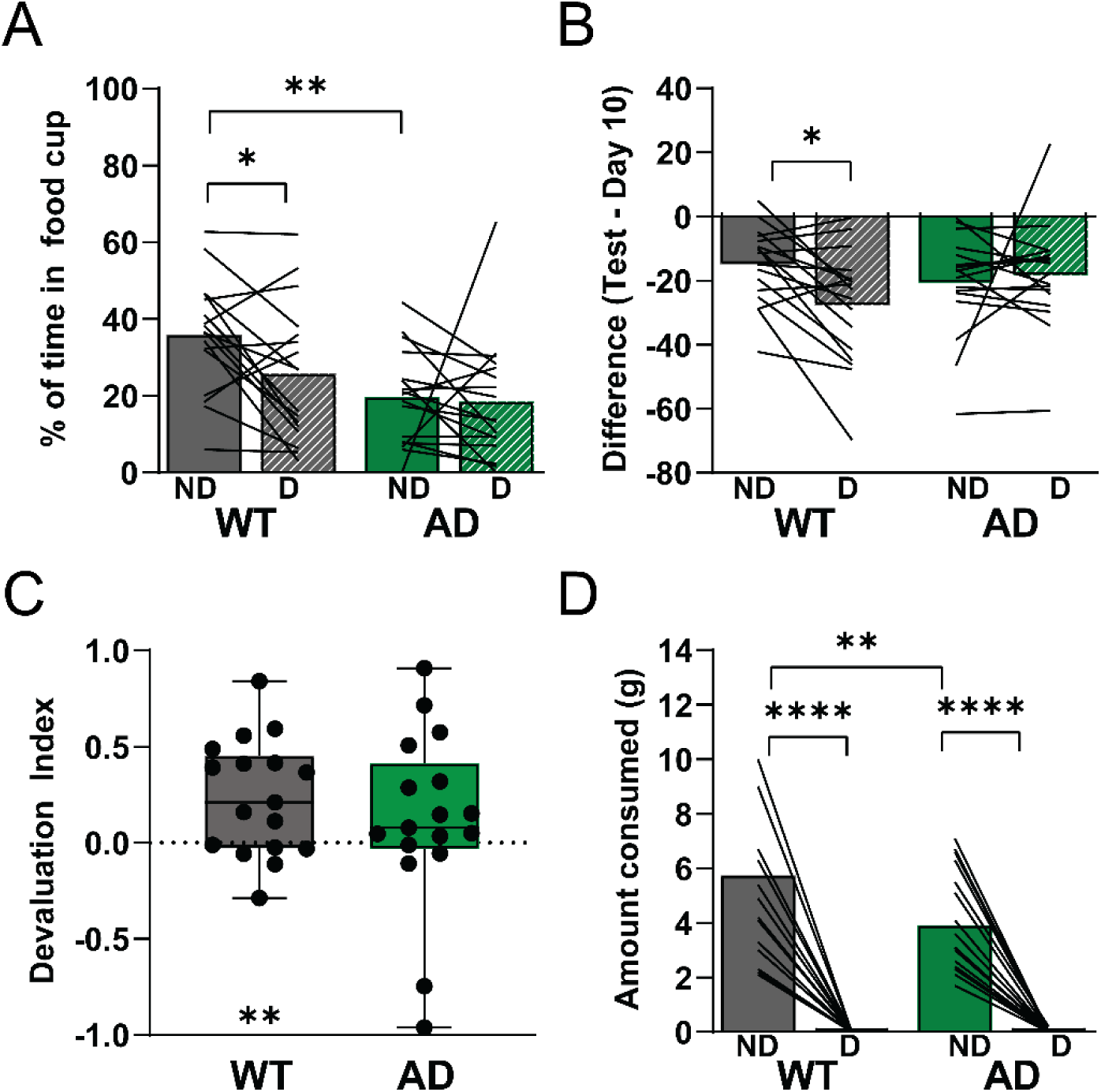
Transgenic AD rats show impaired ability to alter behavior towards reward predictive cues following a devalued expected outcome without impaired consummatory behavior towards the devalued outcome. A) WT rats spent significantly less time in the food cup during the cue that predicted the devalued outcome compared to the cue that predicted the nondevalued outcome, while there was no difference between the time spent in the food cup during the cue that predicted the two outcomes in AD rats [main effect of genotype (F_(1,32)_=8.0, p=0.008, but not of devaluation status (F_(1,32)_=3.0, p=0.091) or devaluation status x genotype interaction(F_(1,32)_=2.05, p=0.16] with Bonferroni planned comparison showing that WT spent more time in the foodcup during cue associated with the nondevalued outcome (*p<0.05), but not the devalued. B) WT animals significantly decreased their responses to cues that predicted the devalued outcome compared to the nondevalued outcome relative to Day 10 behavior, while AD animals did not [no main effect of devaluation status (F_(1,32)_=3.1, p=0.09) or genotype (F_(1,32)_=0.15, p=0.70), a significant devaluation status x genotype interaction effect (F_(1,32)_=6.2, p=0.02)]. Bonferroni post-hoc revealed a significant difference between the change in responding in the devalued condition compared to the nondevalued in WT (* p<0.05), but not in AD rats. C) There was no difference between the WT and AD transgenic rats on the devaluation index (t_32_=0.92, p=0.37), however, only WT rats showed successful flexible behavior as WT showed a significant positive value for the devaluation index, while AD rats’ performance did not differ from 0 indicative of impaired flexible behavior (WT: t_16_=3.2, p=0.005; AD: t_16_=0.32, p=0.33). D) WT and AD rats show successful devaluation in a consummatory test and AD rats also show a decrease in the general consumption of the nondevalued outcome [main effect of devaluation status (F_(1,32)_=126.6, p<0.0001), main effect of genotype (F_(1,32)_=4.5, p=0.04), and interaction between genotype x devaluation status (F_(1,32)_=4.6, p=0.04)]. Bonferroni post-hocs showed that both WT and AD rats ate significantly more of the nondevalued outcome than the devalued outcome (****p<0.001) and that WT rats ate more of the nondevalued outcome, but not the devalued outcome, than AD rats (**p<0.01).

On the post-devaluation test day, we recorded from 280 neurons (AD: 128 WT: 152) in the medial prefrontal cortex (prelimbic and infralimbic subregions). We found that there were three distinct classification of mPFC neurons, neurons that were excited by the cue (increase in cell firing during cue presentation), inhibited by the cue (decrease in cell firing during the cue presentation), or nonphasic (no change in cell firing during the cue presentation) with all neurons recorded from WT and AD rats represented in Figure 6A-B. We found a significant difference in the proportion of neuron classification across groups (AD and WT) and devaluation status (ND and D) as shown by a χ^2^ analysis (χ^2^=15.5, df=6, p=0.02). In order to determine which factor contributed to the overall differing proportions, we examined χ^2^ analysis within WT (ND vs D; χ^2^=1.4, df=2, p=0.49) and AD (ND vs D; χ^2^=0.1, df=2, p=0.95). We did find that when we compared AD vs WT in the nondevalued (χ^2^=4.9, df=2, p=0.08) and the devalued condition (χ^2^=10.3, df=2, p=0.006), there was a significant difference in the devalued condition (a potential trend in the nondevalued). Next, we examined the proportion of only phasic neurons to determine the differences between AD and WT in neural encoding. We found that on the post-devaluation test day that AD rats show an overall greater proportion of neurons that were classified as excited, regardless of the devaluation status (i.e., in both the nondevalued and devalued condition) as shown in Figure 6C (ND: χ^2^=4.9, df=1, p=0.03; (D: χ^2^=8.8, df=1, p=0.003), but no differences in the strength of firing of the individual neurons between group as shown in Figure 6D. Specifically, we calculated an AUC for the cue duration (either nondevalued or devalued) for each neuron and a two-way ANOVA revealed no main effect of group (AD vs WT), devaluation status (nondevalued vs devalued), or interaction for neurons classified as excited (group: F_(1,98)_=1.4, p=0.24; devaluation status: F_(1,98)_=0.01, p=0.93; interaction F_(1,53)_=0.45, p=0.50) or inhibited (group: F_(1,98)_=1.4, p=0.24; devaluation status: F_(1,98)_=0.36, p=0.55; interaction F_(1,98)_=1.8, p=0.18). Thus, there was a greater proportion of neurons that were excited in the AD rats, but these individual neurons did show a greater overall firing changes relative to baseline compared to WT rats.

**Figure 6.**
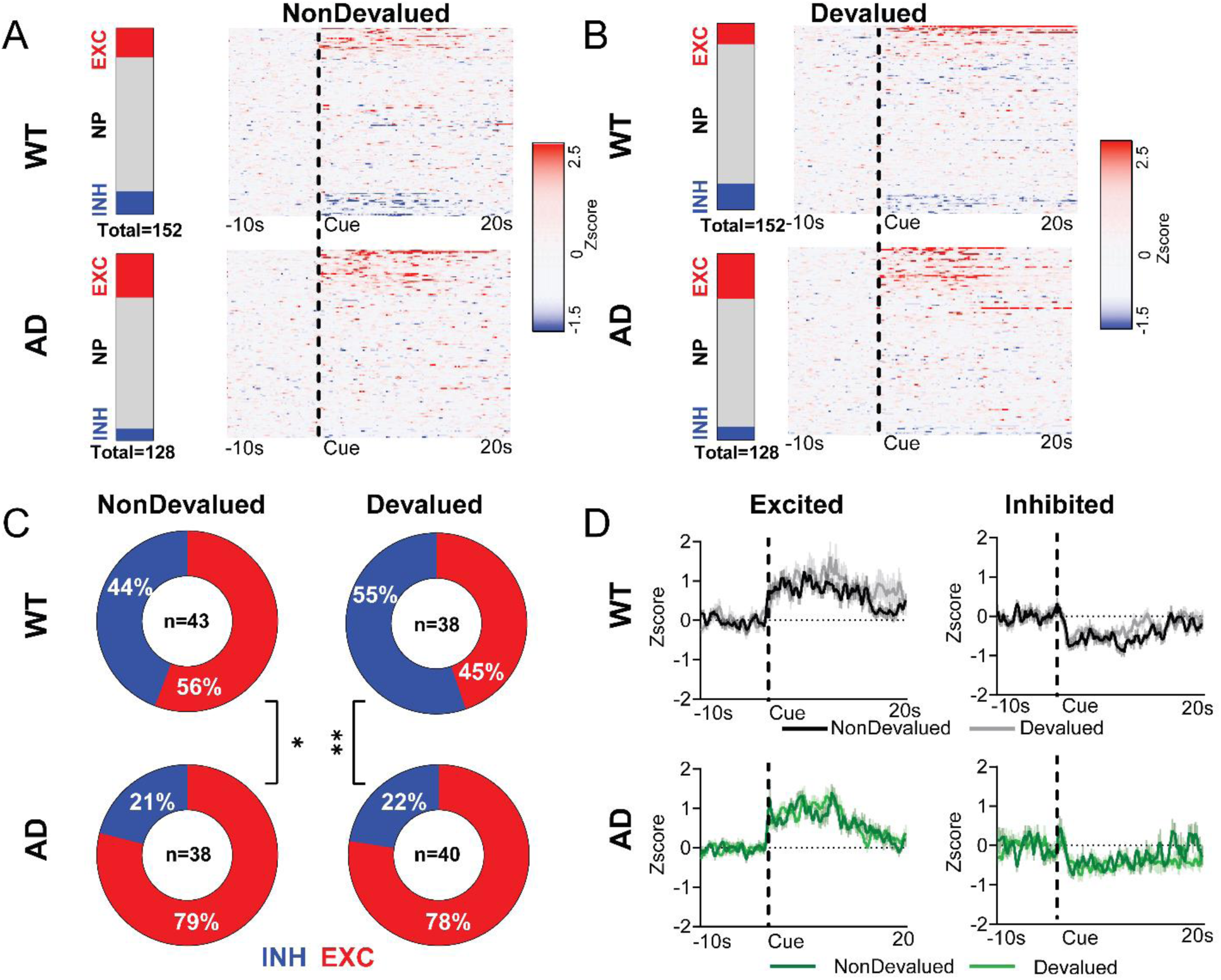
Transgenic rats show hyperactive neural profile compared to WT control during the post-devaluation test session. A) Proportion of excited, inhibited and nonphasic neurons to the cue that predicted the nondevalued outcome in WT and AD and corresponding heatmap of all neurons recorded. B) Proportion of excited, inhibited and nonphasic neurons to the cue that predicted the devalued outcome in WT and AD and corresponding heatmap of all neurons recorded. * represents a significant difference between the proportion of neurons between WT and AD to the devalued cue (χ^2^=10.3, df=2, p=0.006). C) Piecharts representing the proportion of phasic neurons classified as excited (EXC) or inhibited (INH) show a difference in the proportion between AD and WT to cues that predicted either nondevalued (χ^2^=4.9, df=1, * p=0.03) or devalued outcome (χ^2^=8.8, df=1, ** p=0.003). with a greater proportion of excited neurons in both conditions in AD rats. D) The average firing (zscore) across all neurons classified as either excited (left) and inhibited (right) in WT and AD animals show no difference in degree of neural responsiveness between AD and WT animals.

**Figure 7.**
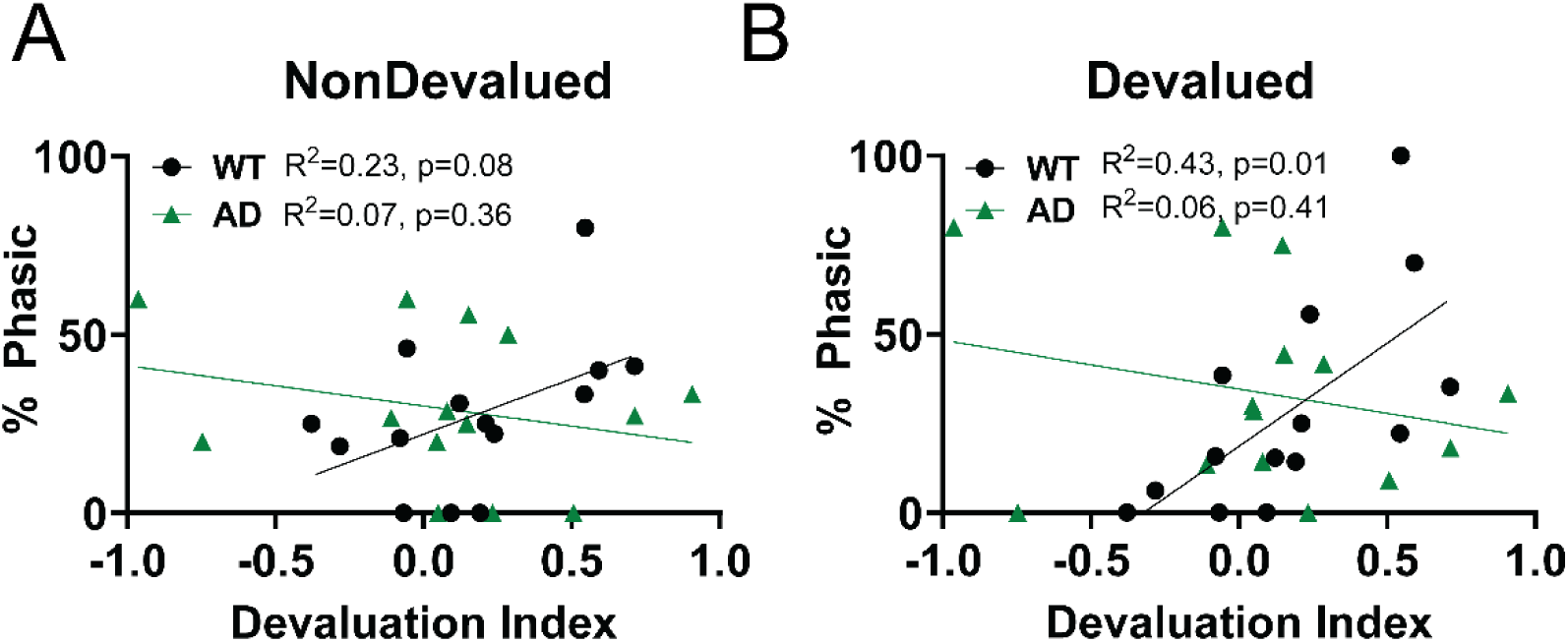
WT rats mPFC neural encoding, but not AD rats, predicts post-devaluation behavior. A) There is no correlation between devaluation index (ND-D/ND+D) and phasic (increase or decrease in cell firing) responsiveness to the cue that previously was paired with the nondevalued outcome, while there is a trend in WT rats. B) There is no correlation between devaluation index (ND- D/ND+D) and phasic (increase or decrease in cell firing) responsiveness to the cue previously paired with the devalued, while there is a significant correlation in that the more phasic responsiveness to the cue predicting the devalued outcome, the better the rats behavioral performance (measured as a devaluation index).

Critically, we examined the correlation between phasic responsiveness (increase and decrease in firing) and the ability to shift behavior post-devaluation in WT (n=14) and AD rats (n=16). We found that WT rats show a significant correlation between the % phasic responding to the cue that predicted the devalued outcome (R^2^=0.43, p=0.01) and a trend between the % phasic responding to the cue that predicted the nondevalued outcome (R^2^= 0.23, p=0.079) and the ability to shift behavior (devaluation index), while there were no correlations between the phasic responding to cues and behavior in AD animals (devalued: R^2^=0.06, p=0.41, nondevalued: R^2^=0.07, p=0.36). Thus, neural activity and flexible behavior are not correlated in AD rats.

## Discussion

We found that AD rats show diminished motivated behavior toward cues that predict rewards, as well as impaired ability to change behavior following a diminished expected outcome. AD rats successfully discriminate between a rewarded cue (CS+) and an unrewarded cue (CS) even though they show diminished approach behavior towards the CS+ (Figure. 2A-C). As such, the deficits observed during conditioning are likely due to diminished motivated behavior. corroborated by the decreased *ab libitum* consumption of the nondevalued outcome in AD rats (Fig. 4D) and consistent with recent work showing diminished motivated behavior towards operant responding (Hernandez et al., 2024; Ostlund et al., 2025). In addition to the overall deficits in motivated behavior in the AD rat, we observe an impaired ability to update behavior towards a devalued outcome. We have previously shown that even at a loss of over 80% of behavioral responding, it is still possible to observe a shift in behavior post-devaluation (West et al., 2012). As such, the loss of overall responding may not account for the inability to change behavior in AD rats. To ensure that the overall decrease in performance was accounted for, we normalized the overall responding to individual animals’ behavior on the last day of conditioning and found that only WT animals showed a difference between the responding in the nondevalued condition versus the devaluation condition (Figure 4B). In addition, only WT animals show an overall positive devaluation index, indicating that they preferentially responded to the cue predicting the nondevalued outcome over the devalued outcome. In contrast AD showed a devaluation index that was not different from 0 indicating that they are responding equally to both cues despite AD rats showing successful discrimination between rewarded and unrewarded cues (Fig. 2C) and successful devaluation of the outcome (Fig 4D). In support of the inflexible AD phenotype, others have shown that AD rats as young as 6 months old are impaired in their ability to shift behavior following reversal in a Morris water maze task (Rorabaugh et al., 2017). In addition, our findings are consistent with impaired flexibility post-devaluation in an instrumental task in 8-9-month-old mouse model of Alzheimer’s disease (Dhungana et al., 2023).

Despite both WT and AD rats showing preferential encoding of the CS+ relative to the CS− (Fig. 3) following conditioning, the proportion of phasic neurons that respond to both events (cue presentation and reward delivery) differs between WT and AD rats (Fig. 4B). Specifically, 52% of WT phasic neurons encode information about both the cue and the outcome, whereas AD phasic neurons are evenly split between cue-outcome (∼1/3), cue-selective (∼1/3), and outcome-selective (∼1/3) encoding. The lack of cue-outcome encoding in AD rats may influence their ability to link the outcome with the cue, and, as a result, disrupt their motivational state towards the cues. In addition, if AD rats do not encode the cue-outcome relationship the same as WT rats during conditioning, the behavioral differences observed on the post-devaluation test day may be a result of AD rats’ inability to integrate the updated outcome value with their previous, potentially weak cue-outcome associations.

In resting state (*i.e.*, anesthetized) TgF344-AD rats, there is general hypofunction in mPFC and hippocampus (Bazzigaluppi et al., 2018; Morrone et al., 2020). However, here we show hyperfunctional mPFC putative pyramidal neuron encoding predictive cues in awake and behaving animals, possibly linked to the behavioral deficits observed following outcome devaluation. In support, increased mPFC activity is associated with worse behavioral performance in reversal learning, suggesting the increased neural activity may drive impaired flexibility (Manning et al., 2019). There is evidence that the excitation/inhibition balance in AD is disrupted (Nataraj et al., 2025). Specifically, local mPFC microcircuits consisting of GABAergic interneurons— particularly parvalbumin-positive interneurons—are dysfunctional in a rodent AD model (Shu et al., 2023), leading to disinhibition of pyramidal projection neurons and hyperexcitability. As such, these changes may be linked to hyperexcitation to cues that we observe in AD animals, and future experiments will aim to determine the mechanism.

Another factor that could be contributing to the pathological mPFC encoding shown in AD rats is alterations to neuromodulatory systems, such as the locus coeruleus-mPFC noradrenergic projection. LC efferents are unique and have distinct functional properties depending on the region of their terminal fields, including the mPFC (Chandler et al., 2014). In humans (Braak and Del Tredici, 2012) and the TgF344-AD rat model (Rorabaugh et al., 2017), the LC is among the first brain regions affected by AD-related neuropathology (tauopathy). In addition, TgF344-AD rats show altered iron homeostasis in the LC, an essential element for general neuronal function (Bhagaloo et al., 2025). Moreover, noradrenergic tone is necessary for flexible behavior as measured by a reversal learning task (Totah et al., 2015) and chemogenetic stimulation of LC rescued reversal-stage reversal learning deficits in 6-month-old AD rats (Rorabaugh et al., 2017). Taken together, these data suggest that a dysfunctional LC-mPFC projection in AD rats could be influencing flexible behavior by disrupting the mPFC encoding and updating of cue-outcome associations. In support, the LC-mPFC projection is necessary for rats’ ability to adaptively respond during a contingency degradation procedure, suggesting that the LC-mPFC projection is key for rats’ ability to update cue-outcome associations (Piccin et al., 2024).

Finally, we found a correlation between mPFC neural activity and the ability shift behavior in WT rats that did not exist in AD rats. Specifically, elevated neural activity during the cue associated with the devalued outcome (and a trend for the cue predicting the nondevalued outcome) predicted enhanced ability to shift away from the devalued outcome. One possibility is that the mPFC becomes activated as rats are integrating information about the cue-outcome associations that require the animals to pay attention, particularly given the role of mPFC in mediating attention (Totah et al., 2013; Bissonette and Roesch, 2015; Del Arco et al., 2017). We have previously found the opposite correlation, in that the greater the elevated prelimbic function, the worse the behavioral performance (Niedringhaus and West, 2022). However, in our prior work, the animals remained engaged with both of the cues longer under extinction than in the current study. It is possible that differential behavioral responses could be due to the use of the Long-Evans strain. As such, distinct behavioral differences between Long-Evans and Fischer rats may underlie some of the behavioral and neural differences observed in the current study and past studies (Niedringhaus and West, 2022). Indeed, we have observed behavioral differences in Long-Evans and Fischer rats in working memory tasks (Gohar et al., 2023) that depend on the medial prefrontal cortex (Auger and Floresco, 2017; Auger et al., 2020; Ciacciarelli et al., 2025).

The mPFC is also functionally active during fear-inducing stimuli (i.e., one that predicts foot shock). Interestingly, animals that behaviorally avoided shock-predictive cues were characterized by a higher proportion of prelimbic neurons that were inhibited during the cue. Conversely, delayed avoidance of the cue was characterized by higher excitation of the prelimbic neurons (Diehl et al., 2018). In further support, photoactivation of the prelimbic, specifically the projection to the nucleus accumbens, impaired avoidance behavior towards a cue predicting shock (Diehl et al., 2020), and the prelimbic to accumbens circuit is critical for behavioral performance in our task (West et al., 2021). As such, it is possible that elevated prelimbic neural activity to a cue that predicts a negative expected outcome may contribute to the impaired change in behavior post-devaluation in AD rats. In support, two distinct phenotypes emerge in rats pressing for a food reward when simultaneously presented with stimuli that predicted shock, i.e., a subset that continued to press for the reward and a subset that suppressed pressing. Rats that continued to press also showed elevated prelimbic neural encoding to the cue during the conflict (Fernandez-Leon et al., 2021), as do rats that continue to respond to (and consume) an outcome that has been devalued (Niedringhaus and West, 2022). Taken together, elevated prelimbic encoding is linked to the continuation of responding to food reward in spite of negative predictive outcomes (either a cue predicting shock or an outcome devalued with conditioned taste aversion).

The ventral (infralimbic, IL) and dorsal (prelimbic, PrL) subregions of the mPFC are canonically considered to exert opposing forces during reward-seeking behavior—the PrL governing goal-directed actions and the IL controlling behavioral inhibition (Killcross and Coutureau, 2003). More specifically, optogenetic inactivation of the IL ablates previously formed habit expression (Coutureau and Killcross, 2003; Smith and Graybiel, 2013), and optogenetic inactivation of the PrL disrupts goal-directed behaviors (Killcross and Coutureau, 2003). However, elevated (Niedringhaus and West, 2022) and depressed (West et al., 2021) PrL activity may be linked to inflexible responding or a loss of value-guided behavior, implicating a potential “inverted U” of PrL activity that does not entirely fit within the framework of traditional theories of PrL/IL functional differentiation. Indeed, both PrL and IL encode contextual information for both the initiation and suppression of behavior (Moorman and Aston-Jones, 2015). In further work regarding mPFC subdivision contribution to natural reward-related behavior, bilateral inactivation manipulations revealed more complexity, including inactivation of both PrL and IL decreased sucrose-seeking behavior during extinction (Caballero et al., 2019). In the current study, we included neurons from both PrL and IL as we observed similar patterns of neural activity to reward predictive cues as others have shown with natural rewards (Moorman and Aston-Jones, 2015).

In summary, our findings demonstrate that AD rats exhibit significant behavioral deficits both in response to reward-predicting cues and in adapting behavior following devaluation of a previously desirable outcome, consistent with their inability to update behavior following outcome shifts and despite intact cue discrimination. The inflexible phenotype is accompanied by disruptions in neural encoding, particularly within the mPFC. Specifically, AD rats fail to display the typical cue-outcome encoding pattern observed in wild-type (WT) rats, which may hinder their ability to integrate updated outcome information, contributing to the observed behavioral impairments. These alterations may contribute to the hyperexcitable mPFC response to reward-predicting cues, exacerbating the deficits in flexible behavior. Furthermore, our data suggest that abnormal mPFC activity is correlated with the failure to shift behavior post-devaluation, highlighting the critical role of mPFC in updating cue-outcome associations. Overall, our findings suggest that deficits in cue-outcome integration and failures to update behavior after outcome devaluation may be driven by disrupted neural encoding in the mPFC, potentially stemming from an altered excitation/inhibition balance and/or neuromodulatory dysfunction. Importantly, our work positions the mPFC as a potential target for therapies aimed at restoring adaptive, value-guided behavior in AD, providing new strategies to combat the core behavioral deficits of the disease.

## Acknowledgments

This work was funded in part by the National Institute on Drug Abuse (R00DA042934 to EAW), the National Institute on Aging (R00DA042934S1, R21AG072355 to EAW), and the NJ Health Foundation (MN), as well as Rowan University School of Osteopathic Medicine internal funds. The authors thank Scott Dunn, Evan Ciacciarelli, Sam Bozarth, and Brianna Linneman for helping maintain our breeding colony and Dr. Ben Rood for assistance with genotyping our animals.

## References

Auger ML, Floresco SB (2017) Prefrontal cortical GABAergic and NMDA glutamatergic regulation of delayed responding. Neuropharmacology 113:10–20.

Auger ML, Meccia J, Phillips AG, Floresco SB (2020) Amelioration of cognitive impairments induced by GABA hypofunction in the male rat prefrontal cortex by direct and indirect dopamine D(1) agonists SKF-81297 and d-Govadine. Neuropharmacology 162:107844.

Bazzigaluppi P, Beckett TL, Koletar MM, Lai AY, Joo IL, Brown ME, Carlen PL, McLaurin J, Stefanovic B (2018) Early-stage attenuation of phase-amplitude coupling in the hippocampus and medial prefrontal cortex in a transgenic rat model of Alzheimer’s disease. J Neurochem 144:669–679.

Beagle AJ, Zahir A, Borzello M, Kayser AS, Hsu M, Miller BL, Kramer JH, Chiong W (2020) Amount and delay insensitivity during intertemporal choice in three neurodegenerative diseases reflects dorsomedial prefrontal atrophy. Cortex 124:54–65.

Berron D, van Westen D, Ossenkoppele R, Strandberg O, Hansson O (2020) Medial temporal lobe connectivity and its associations with cognition in early Alzheimer’s disease. Brain 143:1233–1248.

Bertoux M, O’Callaghan C, Flanagan E, Hodges JR, Hornberger M (2015) Fronto-Striatal Atrophy in Behavioral Variant Frontotemporal Dementia and Alzheimer’s Disease. Front Neurol 6:147.

Bhagaloo KA, Yu L, West EA, Chandler DJ, Shcherbik N (2025) Alterations in iron levels in the locus coeruleus of a transgenic Alzheimer’s disease rat model. Neurosci Lett 850:138151.

Bissonette GB, Roesch MR (2015) Neural correlates of rules and conflict in medial prefrontal cortex during decision and feedback epochs. Front Behav Neurosci 9:266.

Braak H, Del Tredici K (2012) Where, when, and in what form does sporadic Alzheimer’s disease begin? Curr Opin Neurol 25:708–714.

Caballero JP, Scarpa GB, Remage-Healey L, Moorman DE (2019) Differential Effects of Dorsal and Ventral Medial Prefrontal Cortex Inactivation during Natural Reward Seeking, Extinction, and Cue-Induced Reinstatement. eNeuro 6.

Chandler DJ, Gao WJ, Waterhouse BD (2014) Heterogeneous organization of the locus coeruleus projections to prefrontal and motor cortices. Proc Natl Acad Sci U S A 111:6816–6821.

Ciacciarelli EJ, Dunn SD, Gohar T, Joseph Sloand T, Niedringhaus M, West EA (2025) Medial prefrontal cortex to nucleus reuniens circuit is critical for performance in an operant delayed nonmatch to position task. Neurobiol Learn Mem 217:108007.

Cohen RM, Rezai-Zadeh K, Weitz TM, Rentsendorj A, Gate D, Spivak I, Bholat Y, Vasilevko V, Glabe CG, Breunig JJ, Rakic P, Davtyan H, Agadjanyan MG, Kepe V, Barrio JR, Bannykh S, Szekely CA, Pechnick RN, Town T (2013) A transgenic Alzheimer rat with plaques, tau pathology, behavioral impairment, oligomeric abeta, and frank neuronal loss. J Neurosci 33:6245–6256.

Coutureau E, Killcross S (2003) Inactivation of the infralimbic prefrontal cortex reinstates goal-directed responding in overtrained rats. Behav Brain Res 146:167–174.

Creese B, Ismail Z (2022) Mild behavioral impairment: measurement and clinical correlates of a novel marker of preclinical Alzheimer’s disease. Alzheimer’s Research & Therapy 14.

de Siqueira AS, Yokomizo JE, Jacob-Filho W, Yassuda MS, Aprahamian I (2017) Review of Decision-Making in Game Tasks in Elderly Participants with Alzheimer Disease and Mild Cognitive Impairment. Dement Geriatr Cogn Disord 43:81–88.

Del Arco A, Park J, Wood J, Kim Y, Moghaddam B (2017) Adaptive Encoding of Outcome Prediction by Prefrontal Cortex Ensembles Supports Behavioral Flexibility. J Neurosci 37:8363–8373.

Dhungana A, Becchi S, Leake J, Morris G, Avgan N, Balleine BW, Vissel B, Bradfield LA (2023) Goal-Directed Action Is Initially Impaired in a hAPP-J20 Mouse Model of Alzheimer’s Disease. eNeuro 10.

Diehl MM, Bravo-Rivera C, Rodriguez-Romaguera J, Pagan-Rivera PA, Burgos-Robles A, Roman-Ortiz C, Quirk GJ (2018) Active avoidance requires inhibitory signaling in the rodent prelimbic prefrontal cortex. Elife 7.

Diehl MM, Iravedra-Garcia JM, Morán-Sierra J, Rojas-Bowe G, Gonzalez-Diaz FN, Valentín-Valentín VP, Quirk GJ (2020) Divergent projections of the prelimbic cortex bidirectionally regulate active avoidance. eLife 9.

Fernandez-Leon JA, Engelke DS, Aquino-Miranda G, Goodson A, Rasheed MN, Do Monte FH (2021) Neural correlates and determinants of approach–avoidance conflict in the prelimbic prefrontal cortex. eLife 10.

Fowler CF, Goerzen D, Devenyi GA, Madularu D, Chakravarty MM, Near J (2022) Neurochemical and cognitive changes precede structural abnormalities in the TgF344-AD rat model. Brain Commun 4:fcac072.

Freedman M, Oscar-Berman M (1989) Spatial and visual learning deficits in Alzheimer’s and Parkinson’s disease. Brain Cogn 11:114–126.

Gohar T, Ciacciarelli EJ, Dunn SD, West EA (2023) Transient strain differences in an operant delayed non-match to position task. Behav Processes 211:104932.

Hamm V, Heraud C, Cassel JC, Mathis C, Goutagny R (2015) Precocious Alterations of Brain Oscillatory Activity in Alzheimer’s Disease: A Window of Opportunity for Early Diagnosis and Treatment. Front Cell Neurosci 9:491.

Hernandez CM, McCuiston MA, Davis K, Halls Y, Carcamo Dal Zotto JP, Jackson NL, Dobrunz LE, King PH, McMahon LL (2024) In a circuit necessary for cognition and emotional affect, Alzheimer’s-like pathology associates with neuroinflammation, cognitive and motivational deficits in the young adult TgF344-AD rat. Brain Behav Immun Health 39:100798.

Igarashi M, Ma K, Gao F, Kim HW, Rapoport SI, Rao JS (2011) Disturbed choline plasmalogen and phospholipid fatty acid concentrations in Alzheimer’s disease prefrontal cortex. J Alzheimers Dis 24:507–517.

Kaufman LD, Pratt J, Levine B, Black SE (2012) Executive deficits detected in mild Alzheimer’s disease using the antisaccade task. Brain Behav 2:15–21.

Killcross S, Coutureau E (2003) Coordination of actions and habits in the medial prefrontal cortex of rats. Cereb Cortex 13:400–408.

Koenig T, Prichep L, Dierks T, Hubl D, Wahlund LO, John ER, Jelic V (2005) Decreased EEG synchronization in Alzheimer’s disease and mild cognitive impairment. Neurobiol Aging 26:165–171.

Manning EE, Dombrovski AY, Torregrossa MM, Ahmari SE (2019) Impaired instrumental reversal learning is associated with increased medial prefrontal cortex activity in Sapap3 knockout mouse model of compulsive behavior. Neuropsychopharmacology 44:1494–1504.

McNamee D, Liljeholm M, Zika O, O’Doherty JP (2015) Characterizing the Associative Content of Brain Structures Involved in Habitual and Goal-Directed Actions in Humans: A Multivariate fMRI Study. The Journal of Neuroscience 35:3764–3771.

Moorman DE, Aston-Jones G (2015) Prefrontal neurons encode context-based response execution and inhibition in reward seeking and extinction. Proc Natl Acad Sci U S A 112:9472–9477.

Morrone CD, Bazzigaluppi P, Beckett TL, Hill ME, Koletar MM, Stefanovic B, McLaurin J (2020) Regional differences in Alzheimer’s disease pathology confound behavioural rescue after amyloid-beta attenuation. Brain 143:359–373.

Nataraj A, Blahna K, Jezek K (2025) Insights From TgF344-AD, a Double Transgenic Rat Model in Alzheimer’s Disease Research. Physiol Res 74:1–17.

Niedringhaus M, West EA (2022) Prelimbic cortex neural encoding dynamically tracks expected outcome value. Physiol Behav 256:113938.

O’Doherty JP (2011) Contributions of the ventromedial prefrontal cortex to goal-directed action selection. Ann N Y Acad Sci 1239:118–129.

Ostlund SB, Chen G, Kosheleff A, Lueptow LM, Zhuravka I, Frautschy SA, Lam HA, Maidment NT (2025) Early emergence of motivational and hedonic feeding deficits in the TgF344-AD rat model of Alzheimer’s disease. Front Aging Neurosci 17:1572956.

Piccin A, Plat H, Wolff M, Coutureau E (2024) Adaptive Responding to Stimulus-Outcome Associations Requires Noradrenergic Transmission in the Medial Prefrontal Cortex. J Neurosci 44.

Pickens CL, Hougham A, Kim J, Wang C, Leder J, Line C, McDaniel K, Micek L, Miller J, Powell K, Waren O, Brenneman E, Erdley B (2024) Impairments in expression of devaluation in a Pavlovian goal-tracking task, but not a free operant devaluation task, after fentanyl exposure in female rats. Behav Brain Res 458:114761.

Report AsA (2024) 2024 Alzheimer’s disease facts and figures. Alzheimer’s & Dementia.

Rorabaugh JM, Chalermpalanupap T, Botz-Zapp CA, Fu VM, Lembeck NA, Cohen RM, Weinshenker D (2017) Chemogenetic locus coeruleus activation restores reversal learning in a rat model of Alzheimer’s disease. Brain 140:3023–3038.

Shu S, Xu SY, Ye L, Liu Y, Cao X, Jia JQ, Bian HJ, Liu Y, Zhu XL, Xu Y (2023) Prefrontal parvalbumin interneurons deficits mediate early emotional dysfunction in Alzheimer’s disease. Neuropsychopharmacology 48:391–401.

Smith KS, Graybiel AM (2013) Using optogenetics to study habits. Brain Res 1511:102–114.

Sood A, Richard JM (2023) Sex-biased effects of outcome devaluation by sensory-specific satiety on Pavlovian-conditioned behavior. Front Behav Neurosci 17:1259003.

Totah NK, Jackson ME, Moghaddam B (2013) Preparatory attention relies on dynamic interactions between prelimbic cortex and anterior cingulate cortex. Cereb Cortex 23:729–738.

Totah NK, Logothetis NK, Eschenko O (2015) Atomoxetine accelerates attentional set shifting without affecting learning rate in the rat. Psychopharmacology (Berl) 232:3697–3707.

Visser PJ, Verhey FR, Hofman PA, Scheltens P, Jolles J (2002) Medial temporal lobe atrophy predicts Alzheimer’s disease in patients with minor cognitive impairment. J Neurol Neurosurg Psychiatry 72:491–497.

West EA, Carelli RM (2016) Nucleus Accumbens Core and Shell Differentially Encode Reward-Associated Cues after Reinforcer Devaluation. J Neurosci 36:1128–1139.

West EA, Forcelli PA, McCue DL, Malkova L (2013) Differential effects of serotonin-specific and excitotoxic lesions of OFC on conditioned reinforcer devaluation and extinction in rats. Behav Brain Res 246:10–14.

West EA, Forcelli PA, Murnen AT, McCue DL, Gale K, Malkova L (2012) Transient inactivation of basolateral amygdala during selective satiation disrupts reinforcer devaluation in rats. Behav Neurosci 126:563–574.

West EA, Niedringhaus M, Ortega HK, Haake RM, Frohlich F, Carelli RM (2021) Noninvasive Brain Stimulation Rescues Cocaine-Induced Prefrontal Hypoactivity and Restores Flexible Behavior. Biol Psychiatry 89:1001–1011.

